# Transcriptional Landscape of Hepatocellular Carcinoma Reveals that Patient Ethnic-Origin Influences Patterns of Expression

**DOI:** 10.1101/2020.12.01.404285

**Authors:** Rachel Zayas, Artemio Sisson, Ariana Kuhnsman, Bolni Marius Nagalo, Lewis R. Roberts, Kenneth Buetow

## Abstract

The global incidence of hepatocellular carcinoma (HCC) has increased threefold in the last 30 years. In the United States, individuals with ancestry from Asia, Africa and Latin America have a significantly higher risk of developing HCC. However, the molecular mechanisms by which HCC disparities occur remain mostly understudied. Herein, we employed advanced bioinformatics analysis tools to identify genomic drivers that could explain the differences seen among HCC patients of distinct ethnicities (geographic origins). Data from TCGA and open-source software tools HiSTAT, StringTie, and Ballgown were used to map next-generation sequencing (NGS) reads from DNA and RNA, assemble transcripts, and quantify gene abundance. Differential genes/transcripts were mapped to known biomarkers and targets of systemic HCC therapeutics. Four overlapping transcripts were identified between each ethnicity group: FCN2, FCN3, COLEC10, and GDF2. However, we also found that multiple genes are expressed in an ethnicity-specific manner. Our models also revealed that both current and emerging biomarkers fail to capture heterogeneity between patients of different ethnicities. Finally, we have determined that first-line treatment, such as Sorafenib, may be better suited for Asian patients, while Lenvatinib may exhibit better efficacy for Caucasian patients. In conclusion, we have outlined that the pathways involved in early hepatocarcinogenesis may occur in an ethnicity-specific manner and that these distinct phenotypes should be taken into account for biomarker and therapeutic development.

## Introduction

The death rate from hepatocellular carcinoma (HCC), the most common primary liver cancer, has increased by 60% globally in the last 30 years, causes more than 800,000 deaths worldwide, and is the sixth leading cause of cancer-related mortality in the US.^1–5^ HCC is associated with various etiologies such as viral hepatitis, metabolic syndrome, chronic cirrhosis, and cytogenetic hepatitis.^6–8^ Nonalcoholic fatty liver (NAFLD) is a condition of excessive fat accumulation in patients’ liver without alcohol abuse and is associated with obesity and metabolic syndrome. NAFLD can cause various liver diseases, including nonalcoholic steatohepatitis (NASH), which can lead to chronic cirrhosis, and ultimately HCC.^8–11^ NAFLD/NASH associated HCC will soon become the leading cause of HCC in the US, as there are an estimated 20 million patients living with NASH.^8,9,12,13^ Chronic hepatitis B virus (HBV) infection is also a common cause of HCC, especially among US-born African Americans, and foreign-born Asians and Africans.^3,8,9,14^ Although diverse in underlying causes of hepatocarcinogenesis, the prevalence of each risk factor of HCC is shifting in the US with an increase of NASH and this reflects a need to reevaluate if early non-invasive detection methods are suitable to address these changes.

HCC is known to disproportionately affect individuals of African, Asian, and Latin American countries, mainly due to lack of resources and inadequate vaccination programs in these regions.^15–18^ Yet high rates of HCC are not only imposed from these external conditions. HCC also affects African, Native American, Asian and Hispanic born Americans at disproportionate rates with more aggressive phenotypes as compared to Caucasians.^6,7,14^ For example, the overall incidence of HCC in the US is 9.4 (individuals per 100,000 persons), but when adjusted by ethnicity, incidence doubles to 18.6 in Asian patients, 15.7 in African American patients, 11.8 in Hispanic patients, and 7.0 in non-Hispanic Caucasian patients.^19^

It is established that ethnicity can sometimes contribute to heterogeneity in HCC; however, the mechanisms involved in these distinct patient populations remain unknown.^20^ Current hypotheses are that earlier onset and more aggressive phenotypes result from factors such as exposure to environmental toxins in the developing world, viral genotype, and host genomics, but none of these hypotheses have been adequately examined to elucidate how these mechanisms affect the development of HCC in different populations. The frequency of hepatic steatosis (NAFLD) varies between ethnicity groups; genome-wide association studies (GWAS) have found that gene polymorphisms in adiponutrin 3 (PNPLA3) have been associated with NAFLD in African and Latin Americans.^21–23^ Heritability of hepatic steatosis in African and Latin Americans have been reported in several alleles, including missense variants in PNLPA3 and GCKR (Glucokinase Regulator), which may be functional and predispose patients to a higher risk of HCC.^24^ Although differences in HCC between ethnicity groups has been well documented in the literature, there is little analysis on how gene expression contributes to HCC in these patients, which may affect how biomarkers are presented in each patient population. We, therefore, hypothesized that biomarkers would elucidate distinct outcomes in various populations, and therefore certain biomarkers may be better suited for patients of specific geographic (ethnic) origins. Moreover, in alignment with this rationale, we were also curious to determine if targets of systemic treatments should be considered in an ethnicity-specific manner.

The advancements to biomarker and treatment development have not adequately focused on the genomic variation of HCC. Therefore, a more accurate understanding of gene expression between diverse populations will better elucidate the genomic landscape to develop more appropriate diagnostic methods and treatment plans, leading to better patient outcomes. RNA-sequencing (RNA-Seq) allows for high throughput analysis of the complete transcriptome, including coding regions, splice sites, rare events (such as fusion events), and assessment of upregulated or downregulated genes.^32^ To assess the transcriptional landscape, open-source computational tools can be used to determine differential transcript expression (DTE) between healthy liver tissue and HCC tissue samples. These tools map NGS reads from DNA and RNA, assemble transcripts, quantify gene abundance, and then provide statistical modeling for visualization and exploratory analysis of results. Using tumor resected samples from the TCGA, and by combining analysis of RNA-seq data and novel information derived from the interactions of the transcriptome and gene-gene interactions, we have sought to identify distinct transcriptional patterns between diverse patient populations of African, Asian, and Caucasian ethnicities that can be associated with clinical outcomes, early onset of HCC, and disparate phenotypes.

## METHODS

### NIH and CGC (Cancer Genomics Cloud)

Our aim was to assess the transcriptomic landscape of Stage I HCC in African American, Asian and Caucasian liver resected samples. To obtain the Stage I sample ID’s, we filtered the dataset in CbioPortal for Neoplasm Disease Stage I as designed by the American Joint Committee on Cancer (AJCC). In the CGC, we selected the most recent run for African American, Asian, and Caucasian samples for HISAT2, StringTie, and paired FASTQ samples. Upon selecting the data, we chose to run the most current version of the CGC application, Version 1 (Supplemental Figure 1).

### RNA-Seq

RNA-seq raw sequencing reads in .bam format from the TCGA LIHC dataset were filtered for AJCC Neoplasm Disease Stage I to identify genomic differences between people of different ethnicities. Asian (N = 80), African American (N = 8), and Caucasian (N =68) samples were converted to FASTQ paired-end read format using SAM tools FASTQ and compressed to FASTQ.GZ format using CGC Seven Bridges Genomics compress applications to ensure efficient alignment. FASTQ.GZ files were paired by sample ID and aligned using the CGC HISAT2-StringTie application for each ethnicity.

Differentially expressed transcripts, measured by log fold change (FC) for each ethnicity, were determined by a fold-change cutoff of 2.0 and q-value < 0.05. Transcripts that met those thresholds for differential expression were overlapped to observe similarities and differences between each group. Transcripts from Ballgown were then filtered for emerging biomarkers considered relevant to HCC, including osteopontin (SPP1), Golgi Membrane Protein (GOLM1), midkine (MDK), Dickkopf-1 (DKK1), glycoprotein-3 (GPC3), coagulation factor II (F2), Decapping Enzyme Scavenger (DCPS) and alpha-1-fucosidase (FUCA1).^25–28^ Biomarkers included common miRNA and miRNA targets to discern ethnicity differences in the literature. In a separate analysis, common HCC treatments and gene targets associated with their pathways were observed for ethnicity group differences. Treatment targets were then filtered for systemic targets including, VEGFR, PDGFR, RAF, c-MET, FGFR, RET, KIT, Tie2, AXL, and PD-1.^29–31^

### Cytoscape

Ballgown output tables for each ethnicity were further clarified by functional enrichment of differentially expressed genes. Differentially expressed protein-coding genes (DEGs) were identified by filtering Ballgown transcript output tables by q-value (q < 0.05) and logFC (with the cutoff of 1.0 and 2.0). Functional enrichment of these DEGs was obtained using CytoScape Version 3.7.2.^33^ Protein-coding genes were submitted to the Search Tool for the Retrieval of Interacting Gene/Proteins (STRING) application to quantify protein-protein network interactions.^34^ STRING interaction networks were then analyzed for significant Gene Ontology (GO) terms to identify any geographic origin specific interactions. Protein-protein interaction (PPI) networks were generated using the STRING default confidence score cutoff of 0.4, with 0 additional interactors to only visualize the log fold change of the transcripts submitted to STRING (Figure 2). Separate lists of transcripts with a log fold change greater than 2.0 and 1.0 were submitted, respectively.

Transcript lists with a q-value <0.05 and logFC > 1.0 were submitted to STRING for the Asian and Caucasian groups. For the African Americans group, q-value <0.1 and LogFC > 1.0 were used for filtering DEGs. Within Cytoscape for the African American population, the confidence level cutoff was set to the default of 4.0, with 50 additional interactors, as the list generated from Ballgown did not contain enough samples for sufficient statistical power. Interactors are defined as genes/transcripts that were not differentially expressed, but are directly involved in pathways of genes that were differentially expressed. The largest clusters containing more than 2 genes were then submitted to the Cytoscape application Markov Clustering Algorithm (MCL), a clustering algorithm applied to complex biological networks that we used to classify genes based on Gene Ontology.

Networks from each geographic origin were combined into one dataset, with critical identifiers for each shared or not shared, to visualize the gene network pathway relationships. Tables were then submitted to the NCI-Nature Pathway interaction database to identify further, after overlapping gene tables in Rstudio, four genes met the threshold; COLEC10, FCN2, FCN3, and GDF2. A Cytoscape network diagram was generated using the default confidence score cutoff of 0.4 and 30 additional interactors to visualize these “hub” genes’ impact (Figure 2).

### GTEx Database

The Genotype-Tissue Expression (GTEx) is an open-source database to study tissue-specific gene expression in healthy samples (https://www.gtexportal.org/home/). Samples in the GTEx database were collected from 1000 individuals from 54 non-diseased tissue sites using molecular assays such as RNA-Seq, Whole Genome Sequencing, and Whole Exome Sequencing. There were 226 healthy liver samples assessed using the GTEx Multiple Gene Query. Gene clusters were submitted, and a heatmap of healthy tissue genotypes was constructed by comparing liver specific genotypes to multiple other tissue-specific genotypes. Results were reported in transcripts per million (TMP). This heatmap was compared against results reported in the CGC to further assess if differentially expressed transcripts found in HCC were liver-specific.

### Statistical Analysis

The Ballgown (Version 2.8.4) application on the CGC was used for quantifying differential expression by measurement of FPKM measurement. From the TCGA-LIHC dataset, patients denoted as Stage I were selected for analysis. Each ethnicity was put into two groups, normal and HCC, and these groups were provided as inputs to the Ballgown application. For calculation of principal component analysis (PCA), log FPKM at the transcript level was used, and only transcripts that passed initial filtering steps were included in PCA. For all ethnicities, only the first two principal components, PC1 and PC2, were selected as they accounted for the greatest variance between samples (African American PC1: 21.2%, PC2: 15.7%; Caucasian PC1: 15.5%, PC2: 9.9%; Asian PC1: 13.9%, PC2: 10%). A Q-value cutoff of 0.05 was used to analyze differentially expressed genes, with African Americans containing nine transcripts, Caucasian containing 504 transcripts, and Asians containing 242 transcripts. In addition, we also controlled for gender.

## RESULTS

### Overlapping Transcripts Between Ethnicity Groups

A total of 4 transcripts including FCN2, FCN3, COLEC10, and GDF2, were found to be differentially expressed between all three groups with HCC as compared to healthy liver controls (Table 2). Both FCN2 (ficolin-2/liver ficolin) and FCN3 (ficolin-3) transcripts were upregulated in each group; however, the degree of upregulation varied widely. For example, the average fold-change (FC) was 3.92, 2.60 and 2.20 for African American, Asian and Caucasian patients, respectively (Figure 1). COLEC10 (collectin superfamily member 10) was upregulated in all ethnicity groups showing heterogeneous results between groups with a FC of 3.25, 2.83 and 1.42 for Asian, African American and Caucasian patients, respectively. Growth differentiation factor 2 (GDF2) had the most similar upregulation between all three groups, comprising a FC of 3.00 for Asian patients, 2.52 for African American patients, and 2.38 for Caucasian patients.

**Figure 1.**
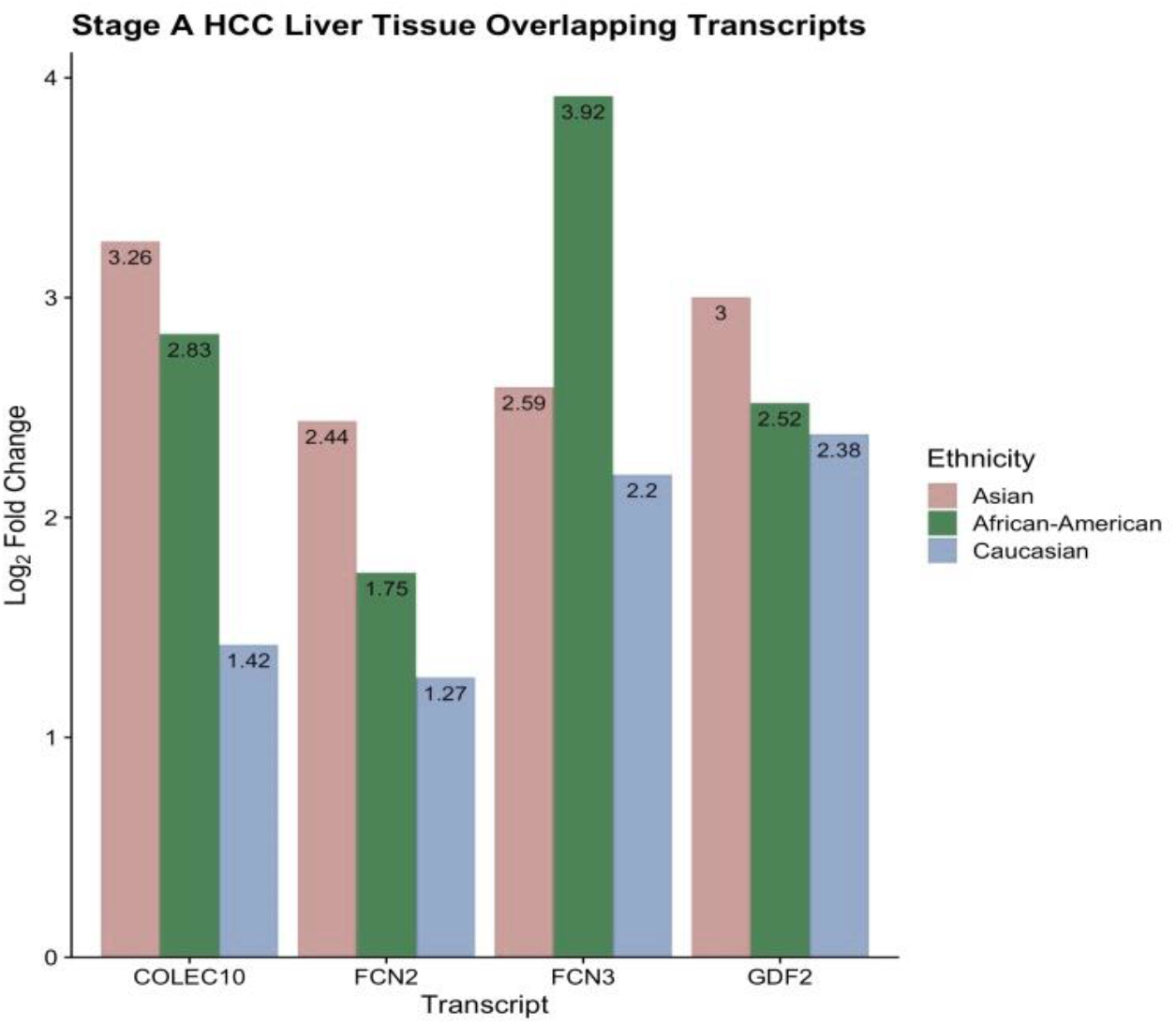
The top most differentially expressed genes overlapping between Asian, African American and Caucasian patient populations (sorted by absolute log_2_ fold change, q-values <0.05, p-value <0.001). Four genes were found to be differentially expressed between all 3 ethnicity groups; GDF2, FCN2, FCN3 and COLEC10. Abbreviations: Growth Differentiation Factor 2 (GDF2), Ficolin-2 (FCN2), Ficolin-3 (FCN3), Collectin Superfamily member 10 (COLEC10).

**Figure 2.**
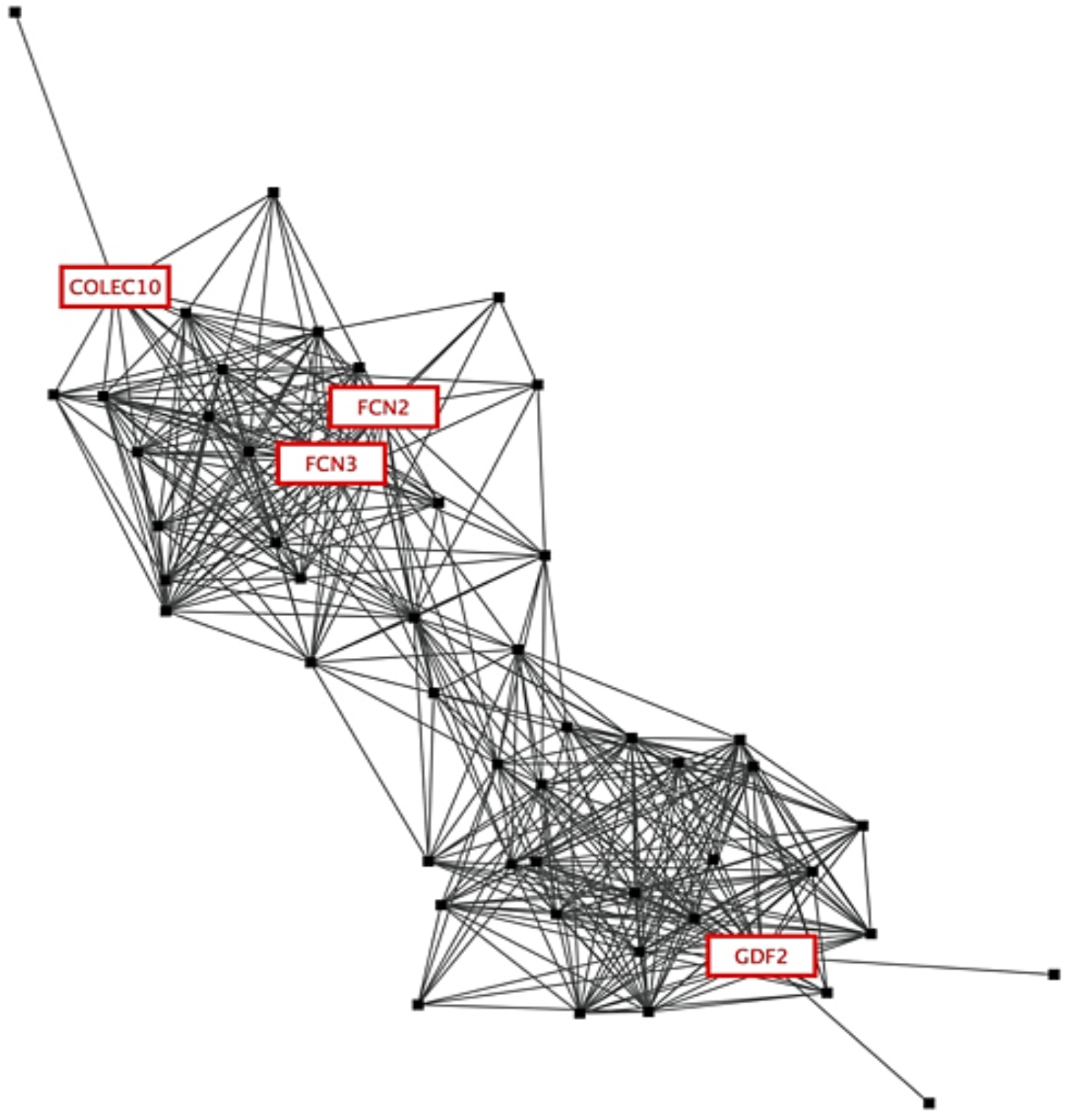
STRING protein-protein interaction (PPI) Cytoscape network. Overlapping differentially expressed transcripts were submitted to STRING with a confidence cutoff of 0.4 and 50 additional interactors (denoted by black squares). Cytoscape application clusterMaker2 MCL cluster was used for relational clustering that describes the network the transcripts are present across all 3 ethnicities.

### Ethnicity Distinctions in Transcriptional Landscape

Although several transcripts showed similar expression patterns between groups, the distinctions in specific transcripts indicated that pathways may be dysregulated in an ethnicity specific-manner. African American patients had 3 exclusively expressed transcripts. In Caucasian and Asian groups, there were more uniquely expressed transcripts within each group than between any groups (Supplemental Figure 2). For example, Asian patients had 134 transcripts that were exclusively expressed in this group only, and Caucasian patients had 176. In contrast, there were 37 overlapping transcripts between Asian and Caucasian patients, 0 overlapping transcripts between African American and Caucasian patients, and 0 overlapping transcripts between African American and Asian patients.

We calculated the representation factor (defined as the number of overlapping genes divided by expected value) and the probably of gene overlaps occurring between two independent datasets (http://nemates.org/MA/progs/representation.stats.html). Between African American (7 transcripts) and Asian (175 transcripts) there were 4 overlapping transcripts, which yielded a representation factor of 98.0, probability (p) < 3.863e-08. Between African American (7 transcripts) and Caucasian (217 transcripts) there were 4 overlapping transcripts, which yielded a representation factor of 79.0, p<9.163e-08. Between Caucasian (217 transcripts) and Asian (175 transcripts) there were 37 overlapping transcripts and yielded a representation factor of 32.4, p< 2.367e-50.

For Caucasian patients, we found that the MT1H (metallothioneins 1H) transcript was the most differentially expressed upregulated gene in Caucasian samples and was not expressed in any other group (Supplemental Figure 3). Additionally, 27 MT (metallothioneins) transcripts and isoforms were significantly upregulated exclusive to Caucasian samples only (data not shown). A full list of up- and downregulated genes for each group can be found in Supplemental Table 1. For Asian patients, the 10 most significant upregulated transcripts included, GNE, SLC38A2, SDS, LSR, AZGP1P1, ADAR, CPN1, APH1A, SORD2P and RPL39P3. For African American patients, the top differentially expressed transcripts included WRN1P1, MYD88, and RDH16 (which were exclusively expressed in this group only) and no transcripts were found downregulated. In our data set, tumor samples showed WRN1P1 upregulation of 3.55 fold-change compared to healthy controls. WRN1P1 has not previously been reported in HCC.

**Table 1.** HCC Demographics including gender, vital status, age, AFP at procurement, vascular invasion, AJCC Neoplasm Disease Stage Classification, fibrosis score, ablation therapy, bilirubin total, etiology and Child Pugh Classification. 100% of patients assessed had Stage I Neoplasm Disease Stage at time of diagnosis as reported by the AJCC. Abbreviation: alpha fetoprotein, AFP. nonalcoholic fatty liver disease, NAFLD, Hepatitis C Virus, HCV, Hepatitis B Virus, HBV, American Joint Committee on Cancer (AJCC).

| **Ethnicity** | **Asian** | **African**<br>**American** | **Caucasian** |
| --- | --- | --- | --- |
| Total Patients | n = 80 | n = 8 | n=68 |
| Male | 62 (78%) | 7 (88) | 43 (63%) |
| Female | 18 (22%) | 1 (12%) | 25 (37%) |
| Vital Status |  |  |  |
| Alive | 70 (88%) | 5 (62%) | 43 (63%) |
| Deceased | 10 (12%) | 3 (38%) | 25 (37%) |
| Age at Diagnosis |  |  |  |
| <40 | 7 | 0 | 3 |
| 41-50 | 16 | 0 | 4 |
| 51-60 | 27 | 1 | 15 |
| 61-70 | 22 | 6 | 25 |
| > 70 | 8 | 1 | 21 |
| AFP at Procurement |  |  |  |
| <400 ng/mL | 58 (72.5) | 7 (88%) | 44 (65%) |
| >400 ng/mL | 20 (25%) | 1 (12%) | 7 (10%) |
| NA | 2 (2.5%) |  | 17 (25%) |
| Vascular Invasion |  |  |  |
| Macro | 62 (77.5%) |  | 1 (1%) |
| Micro | 14 (17.5%) | 1 (12%) | 3 (4%) |
| None | 4 (5%) | 7 (88%) | 52 (77%) |
| NA | 0 |  | 12 (18%) |
| Stage |  |  |  |
| I | 80 (100%) | 8 (100%) | 68 (100%) |
| Liver Fibrosis |  |  |  |
| No Fibrosis | 6 (7.5%) | 2 (25%) | 21 (31%) |
| Portal Fibrosis | 12 (15%) | 1 (12.5%) | 4 (6%) |
| Fibrous Speta | 8 (10%) | - | 5 (7%) |
| Nodular Formation | 1 (1.3%) | - |  |
| Established Cirrhosis | 29 (36.3%) | 1 (12.5%) | 12 (18%) |
| NA | 24 (30%) | 4 (50%) | 26 (38%) |
| Ablation Therapy |  |  |  |
| Yes | 4 (5%) | - | 3 (4%) |
| No | 69 (86.3%) | 3 (37.5%) | 36 (53%) |
| NA | 7 (8.8%) | 5 (62.5%) | 29 (43%) |
| Bilirubin Total |  |  |  |
| <0.5 | 21 (26%) | 2 (12.5%) | 30 (44%) |
| >0.5 | 59 (74%) | 4 (50%) | 28 (41%) |
| NA | - | 2 (12.5%) | 10 (15%) |
| Etiology |  |  |  |
| HBV | 46 (58%) | 0 | 1 (1.5%) |
| Alcohol Consumption | 4 (5%) | 0 | 17 (25%) |
| No Known Risk Factor | 4 (5%) | 1 (12.5%) | 20 (29%) |
| NAFLD | 1 (1%) | 0 | 3 (4%) |
| HCV | 4 (4%) | 3 (37.5%) | 8 (12%) |
| N/A | 4 (5%) | 1 (12.5%) | 5 (7%) |
| Two or more factors | 17 (21%) | 3 (37.5%) | 13 (19%) |
| Hemochromatosis | 0 | 0 | 1 (1.5%) |
| Child Pugh Classifications |  |  |  |
| A | 76 (95%) | 5 (62.5%) | 37 (54%) |
| B | 3 (4%) | 1 (12.5%) | 3 (4%) |
| C | 0 | 0 | 1 (1.5%) |
| N/A | 1 (1%) | 2 (25%) | 27 (40%) |

**Table 2.** Description of overlapping genes found between each patient populations of different ethnicities. Function of each gene in the liver is described. Abbreviations: Growth Differentiation Factor 2 (GDF2), Bone Morphogenetic Protein 9 (BMP9), Collectin Subfamily Member 10 (COLEC10), CL-L1 (Collectin Liver 1), Ficolin 2/Fibrinogen Domain Containing Protein 2 (FCN2), Ficolin-3/Fibrogen Domain Containing Protein-3 (FCN3).^38–43^

| **Gene** | **Function in the Liver** |
| --- | --- |
| **GDF2** | Primarily expressed in the liver and associated with liver fibrosis through Smad signaling. |
| **COLEC10** | Encodes Collectin Liver 1 (CL-L1) part of the lectin pathway complement cascade; generally upregulated in healthy liver. |
| **FCN2** | Ficolin-2 is involved in the lectin complement cascade, is synthesized in the liver and can recognize microbial binding sites in circulation, FCN2 is often downregulated in HCC as compared to healthy controls. |
| **FCN3** | Ficolin-3 also involved in the lectin pathway and activates the complement cascade through opsonic properties, FCN3 is also often downregulated in HCC as compared to healthy controls. |

### Genotype Transcript Expression Database

We then assessed liver specificity for each gene that showed differential expression between groups. We found that FCN2 and FCN3 transcripts were highly expressed in healthy liver tissue, but were also found upregulated in multiple other tissues, including a higher expression in lung tissue (FCN3 > 1100 TMP) then in the liver (FCN3 > 67 TMP). FCN2 and FCN3 transcripts were found with similar gene expression profiles in the prostate, thyroid, spleen, kidneys and heart. GDF2 (BMP9) was expressed primarily in liver tissue, as compared to all other tissues assessed (Figure 3). Additionally, COLEC10 was also primarily expressed in liver tissue as compared to all other tissues reported in this assessment. (A full list of GTEx profiles of genes expressed in an ethnicity-specific manner can be found in Supplemental Figure 4).

**Figure 3.**
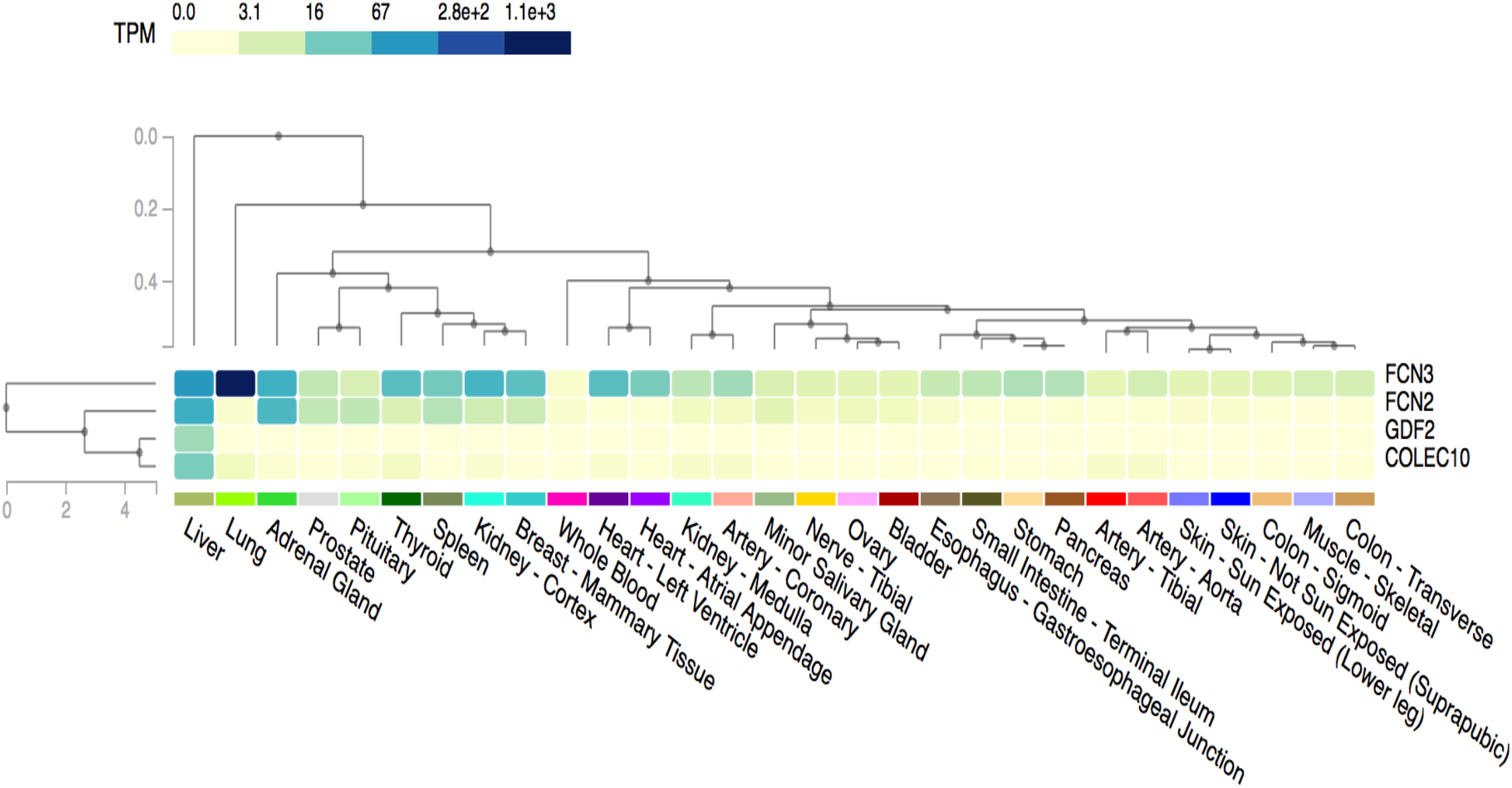
Healthy liver-specific expression patterns of overlapping transcripts found in Stage I HCC. Using the GTEx Database Multiple Gene Query Assessment, overlapping transcripts found between all 3 ethnicities were input into query, heatmap was constructed to assess liver specificity. Map is clustered by tissue. Legend (top left) depicts extent of transcript expression in transcripts per milliohm (TPM).

### Current and Emerging Biomarkers

We assessed our outputs to determine the gene and transcript expression of these biomarkers and various miRNA targets. We found that none (neither combined, nor individual group) had statistical significance within the criteria of absolute value logFC > 2.0, p-value <0.05, and q-value <0.05. We wanted to extrapolate if there were distinctions between patients of different ethnicities, so we expanded the sample size using TCGA data to include all stages of HCC to assess patterns for current and emerging biomarkers. The sample size of all stages of HCC now included 158 Asian patients, 17 African American patients, and 153 Caucasian patients. We followed the same steps outlined in S. Figure 1 as followed for the previous gene expression analysis (patient demographics in Supplemental Table 2).

In our analysis, two biomarkers, GPC3 and MDK, and three miRNAs targets, FSCN1, CD82, and P1M1, were found to be differentially expressed in specific groups with cutoffs of absolute value LogFC > 2.0, p-value <0.01, and q value<0.01 (Table 3). Both GPC3 and MDK were differentially expressed in Caucasian and Asian HCC samples. There was no statistical significance for African American patients for these biomarkers, nor any other biomarkers assessed. In Asian patients, GPC3 had a −3.48 logFC, while Caucasian samples showed a −2.86 logFC. MiRNA targets FSCN1 (Fascin actin-bundling protein 1) and CD82 (CD82 molecule, metastatic tumor suppressor) were found to be differentially expressed in Asian samples only, while miRNA target P1M1 (Pim-1 Proto-Oncogene, Serine/Threonine Kinase) showed significant upregulation in Caucasian samples only (Table 3).

**Table 3.** Biomarker assessed in each ethnicity group. Assessment of differential expression of commonly used biomarkers including midkine (MDK) and glycoprotein-3 (GPC3), and miRNA targets Fascin actin-bundling protein 1 (FSCN1), CD82 molecule/ metastatic tumor suppressor (CD82) and Pim-1 Proto-Oncogene, Serine/Threonine Kinase (P1M1). Biomarkers reported fell within criteria of absolute value LogFC > 1.0, p-value <0.001, and q value <0.002. Blank box indicates non-significant results.

|  | **Asian**<br>**LogFC** | **African American**<br>**LogFC** | **Caucasian**<br>**LogFC** |
| --- | --- | --- | --- |
| **MDK** | -2.41 | - | -2.15 |
| **GPC3** | -3.48 | - | -2.86 |
| **FSCN1** | 1.50 | - | - |
| **CD82** | 1.65 | - | - |
| **P1M1** | - | - | 1.73 |

**Table 4.**
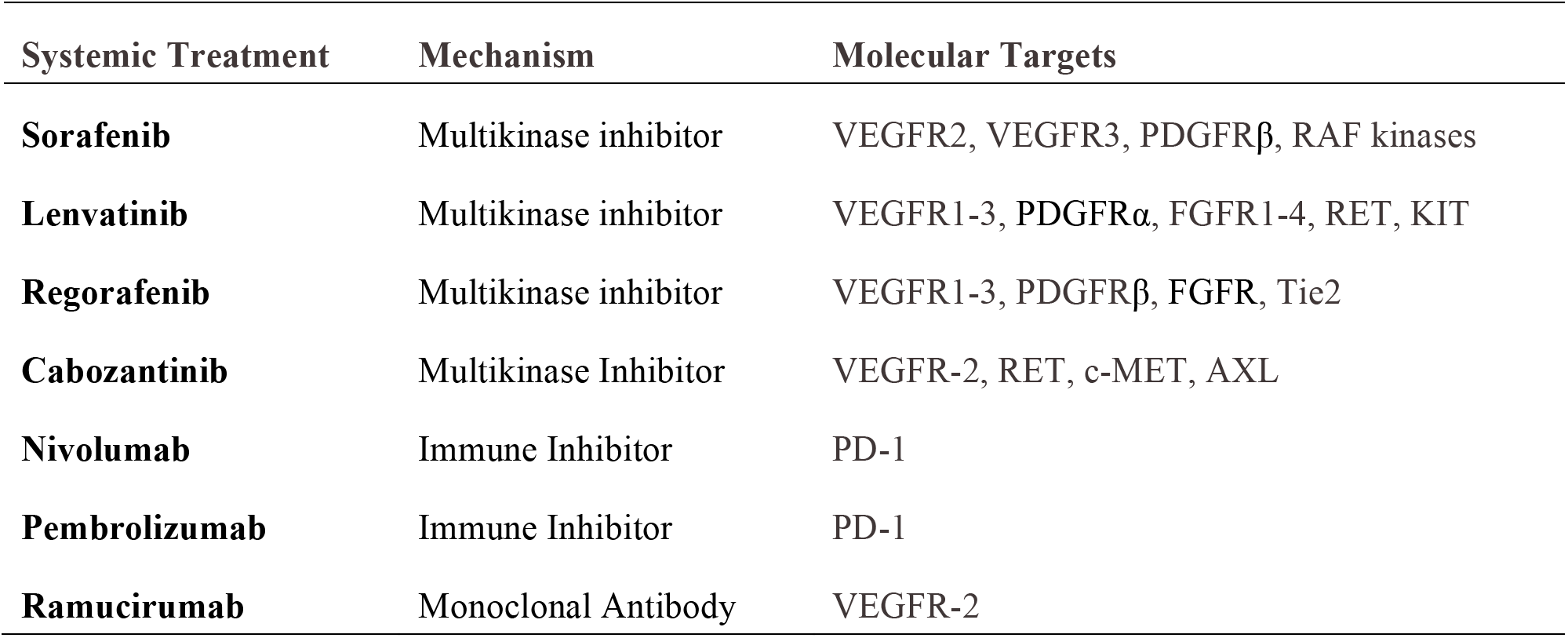
Systemic Treatments for HCC, stratified by treatment name, molecular mechanism and gene targets. Abbreviations: Vascular endothelial Growth Factor Receptor (VEGFR), Fibroblast Growth Factor Receptors (FGFR), Platelet Derived Growth Factor Receptors (PDGFR), Raf-1 Proto-Oncogene or Serine/Threonine Kinase (Raf-1 Kinases), RET Proto-Oncogene/Receptor Tyrosine Kinase (RET), KIT Proto-Oncogene/Receptor Tyrosine Kinase (KIT), TEK Receptor Tyrosine Kinase/Angiopoietin-1 Receptor (Tie2), MET Proto-oncogene/Receptor Tyrosine Kinase/Hepatocyte Growth Factor Receptor (c-MET), AXL Receptor Tyrosine Kinase (AXL) and Programmed Cell Death 1 (PD-1).^29–31^

| **Systemic Treatment** | **Mechanism** | **Molecular Targets** |
| --- | --- | --- |
| **Sorafenib** | Multikinase inhibitor | VEGFR2, VEGFR3, PDGFRβ, RAF kinases |
| **Lenvatinib** | Multikinase inhibitor | VEGFR1-3, PDGFRα, FGFR1-4, RET, KIT |
| **Regorafenib** | Multikinase inhibitor | VEGFR1-3, PDGFRβ, FGFR, Tie2 |
| **Cabozantinib** | Multikinase Inhibitor | VEGFR-2, RET, c-MET, AXL |
| **Nivolumab** | Immune Inhibitor | PD-1 |
| **Pembrolizumab** | Immune Inhibitor | PD-1 |
| **Ramucirumab** | Monoclonal Antibody | VEGFR-2 |

### Current and Emerging Treatments

Similar to the findings with existing and emerging biomarkers, no targets of systemic treatments were found to be differentially expressed in any group with the original sample size. Thus, as outlined above, we assessed all stages of HCC from the TCGA database. No statistically significant results were found for systemic treatment targets for African American patients. MET (a drug target of Cabozantinib) showed downregulation FC of −1.58 in Asian samples compared to FC of −0.76 in Caucasian samples (Table 5). Meanwhile, PDCD5 (target of Nivolumab and Pembrolizumab) showed a downregulation FC of −1.46 in Asian samples and −0.79 in Caucasian samples.

**Table 5.** Transcriptional expression of systemic treatments for HCC stratified by patient ethnicity. Specified gene target reported with systemic treatment name. Reported values fell within criteria of absolute value LogFC >1.0, p-value <0.001, and q value <0.002. Blank box (-) indicates non-significant results. *Indicates results were non-significant, but reported. Absence of molecular target (gene name) indicates results were non-significant.

| **Gene** | **Treatment** | **Asian FC** | **African**<br>**FC\*** | **Caucasian FC** |
| --- | --- | --- | --- | --- |
| **MET** | Cabozantinib | -1.58 | - | -0.76 |
| **ARAF** | Sorafenib | 1.20 | 1.29 | 0.51 |
| **RAF1** | Sorafenib | - | 1.24 | -0.50 |
| **PDGFRα** | Lenvatinib | 1.03 | 1.46 | 0.75 |
| **FGFR2** | Lenvatinib | - | 2.26 | - |
| **FGFRL1** | Lenvatinib / Regorafenib | -1.39 | - | 0.68 |
| **FGFR1OP2** | Lenvatinib / Regorafenib | -1.13 | -0.50 | -0.60 |
| **VEGFR1** | Lenvatinib / Regorafenib | - | 1.25 | -0.63 |
| **PDCD5P1** | Nivolumab /<br>Pembrolizumab | -0.82 | -0.56 | -0.46 |
| **PDCD5** | Nivolumab /<br>Pembrolizumab | -1.46 | -0.79 | -0.79 |

Systemic treatments Lenvatinib and Regorafenib are multikinase inhibitors (that block kinase activity to hinder angiogenesis, metastasis, and oncogenesis) and both target FGFRL1 and FGFR1OP2.^36^ Interestingly, FGFRL1 showed a FC downregulation of −1.39 in Asian patients and an upregulation of 0.68 in Caucasian patients. In accordance with this, FGFR1OP2 showed a FC of −1.13 in Asian patients, compared to FC of −0.60 in Caucasian patients. This may indicate that Lenvatinib and Regorafenib may be better suited for inhibiting Caucasian patients’ tumor environment (based upon an action potential to downregulate target), as Asian patients already exhibited significant downregulation of FGFRL1. In addition, each ethnicity group GO terms consisted of leukocyte mediated immunity, regulation of MAPK cascade and positive regulation of cellular processes (Figure 4). Both Caucasian patients and African American patients reported positive regulation of immune system processes (data not shown).

**Figure 4.**
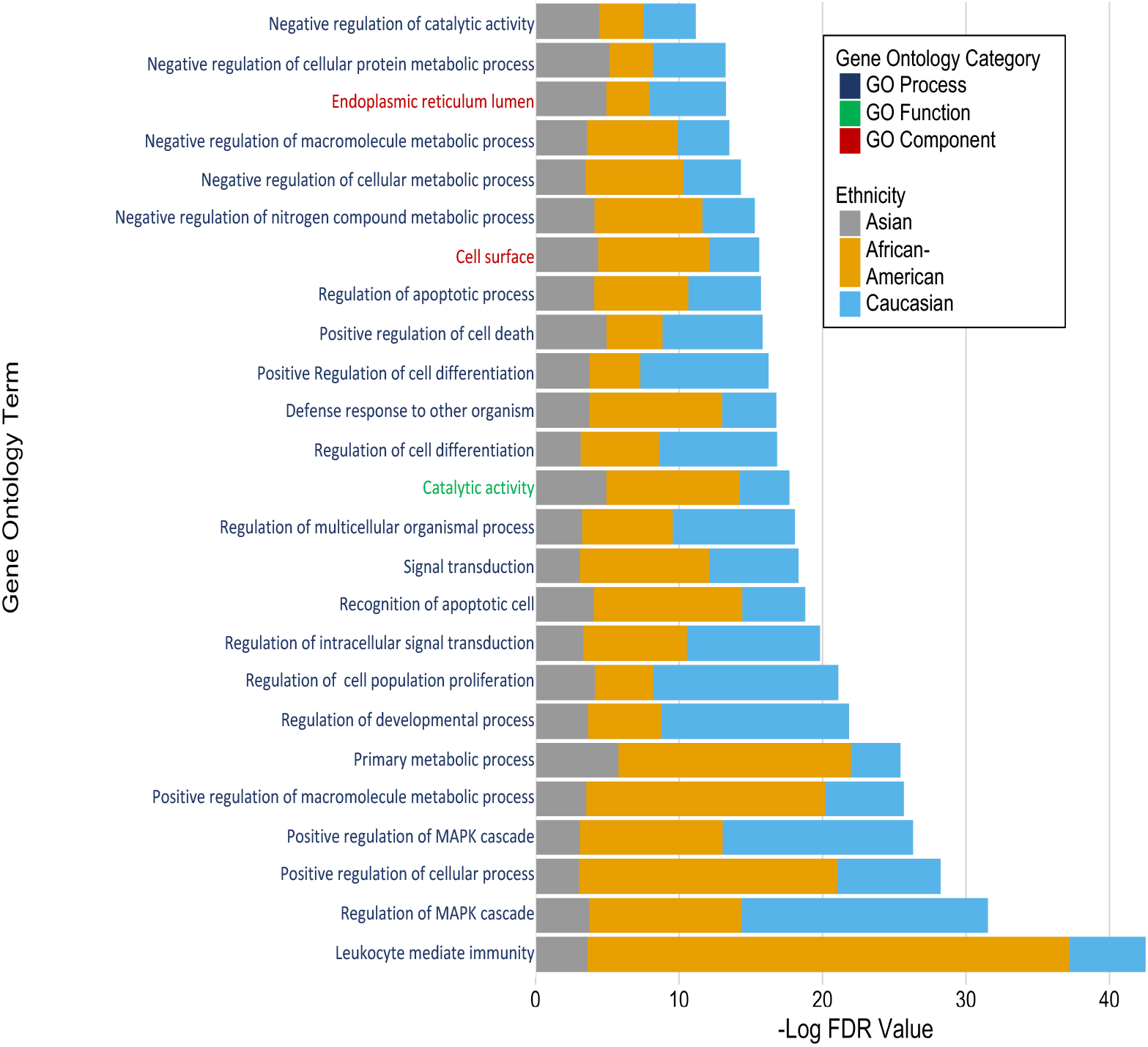
Gene Ontology Bar Plot. Top-25 Gene Ontology (GO) stratified across ethnicity and GO term classification. Differentially expressed transcript lists were submitted to STRING for each ethnicity with a confidence level cutoff of 0.4. Gene Ontology classifications including GO Component, GO Function, and GO Process were obtained from STRING. The mean -Log FDR for each of the terms were calculated and the top-25 highest significance values were visualized. Classification of each GO-term are color-coded on the y-axis, while the -Log FDR value is present on the x-axis.

## DISCUSSION

### Overlapping Transcripts

We conducted a study to decipher transcriptional mechanisms that contribute to heterogeneity in patients of Asian, African and Caucasian ethnicities. We identified four transcripts, FCN2, FCN3, COLEC10, and GDF2 (BMP9), that were differentially expressed during early-stage HCC between the three ethnicity groups assessed. Ficolins are primarily synthesized in the liver and are found in serum as molecules involved in innate immunity that can recognize a variety of microbial binding sites.^38^ Ficolins are involved in the lectin pathway of complement along with two collectins, including mannan-binding lectin (MBL) and Collectin Liver 1 (CL-L1) (Figure 2). In our findings, the COLEC10 transcript, which encodes CL-L1, was found to be upregulated in all three ethnic groups. This suggests that the lectin pathway of complement cascade may be a key indicator of early hepatocarcinogenesis.

In contrast to our findings, FCN2, FCN3, and COLEC10 are known to be downregulated during HCC compared to healthy controls.^38–41^ Nevertheless, FCN2, FCN3 and COLEC10 were each upregulated in our analysis, and the degree of upregulation varied widely. For Caucasian patients, (FCN2 and FCN3) showed the lowest expression; this may reflect Caucasian patients’ poor prognosis in this data set. Specifically, 25 of 68 patients, or 37% of Caucasian patients diagnosed with Stage I in this dataset are now deceased, which is slightly higher than the overall vital status of 24% from this data set (Table 1). Since our findings are in direct contrast to the literature, this may indicate that stage of HCC may be associated with activity of the complement cascade. Further assessment should investigate the lectin pathway involvement and specific microbiome signatures that are active in a stage-specific manner.

In addition to the dysregulation of the complement cascade, GDF2 (BMP9) was also upregulated in all three ethnicity groups. BMP9 is primarily expressed in liver cells, has been associated with fibrosis, and is involved in the PI3K/AKT and p38MAPK cascades.^42,43^ Overexpression of BMP9 has been reported in 40% of HCC tissue, and a recent finding suggests that BMP9 activation in HCC cells must be accompanied by p38MAPK activation to induce and sustain tumorigenic phenotypes in liver cells. This indicates that if BMP9 were to be used in biomarker development, this target should be multiplexed with alternative genes involved in the p38MAPK cascade.

Tissue specificity was considered for each biomarker. FCN2 and FCN3 were found upregulated in multiple other tissues in the body, and since they would not be liver specific if found in circulation, neither would be suitable biomarker candidates. GDF2 and COLEC10 were both found overexpressed in our dataset (Table 2) and found primarily expressed in healthy liver-tissue. Therefore, GDF2 and COLEC10 should be investigated as biomarkers candidates, and expression in circulation should be assessed with the knowledge that transcripts showed distinct expression patterns within each group.

### Differential Transcripts Between Groups

Although there were several overlapping transcripts, many transcripts were expressed in an ethnicity-specific manner. For Caucasian patients, 27 MT transcripts were represented in this group only. MTH1 is a subtype of MT genes, and wild-type expression regulates oxidative damage which can facilitate proliferation when downregulated.^44^ In African American patients, WRN1P1, MYD88, and RDH16 were exclusively expressed in this group only. The mutant version of WRN has been associated with malignant tumors and genomic stability.^46^ In Asian patients, SLC38A2 (Solute Carrier Family 38 Member 2) was uniquely expressed in this group only. SLC38A2 is expressed in almost every tissue and has been reported to be elevated in patients with precancerous lesions associated with viral hepatitis.^45^

While the association between SLC38A2 might be related to the fact that 89% of Asian patients in this study had viral hepatitis as the primary risk-factor (including patients with two or more risk factors) a deeper assessment suggests that microbiome alterations may contribute to early tumorigenesis. For example, MARCO (macrophage receptor with collagenous structure) is often involved in innate antimicrobial responses and was found upregulated in both Asian patients (FC 4.50) and Caucasian patients (FC 3.64) (data not shown).^35^ Nonetheless, only 25% (17 of 68) of Caucasian patients had a primary risk factor of viral hepatitis, suggesting that antimicrobial responses other than viral hepatitis may be involved in progression to HCC. The functional role of the microbiome and discrete pathways should be further explored.

### Biomarkers

Distinctions between current and emerging biomarkers identified that only two biomarkers, GPC3 and MDK, and three miRNAs FSCN1, CD82, and P1M1, were found expressed in Asian and Caucasian patients. Of these, only GPC3 and MDK were found in more than one group (Asian and Caucasian). If geography, ethnic origin, gender, or distinct etiologies present with specific biomarkers, these discrepancies must be incorporated into biomarker development to appropriately assess patients. GPC3 and MDK showed superior results as compared to current alternatives using AFP and other biomarkers on the tissue level (data not shown). Distinct expression patterns of biomarkers in each ethnic group indicate that current biomarkers must consider heterogeneity between groups or HCC phenotypes in general, and current approaches to biomarker development should be reconsidered to better account for the high degree of variability. We recommend a multi-target assessment to address these challenges.

### Systemic Treatment Assessment

Transcriptional expression of drug targets should be interpreted with caution. We identified that for Asian patients, Sorafenib might be the best first-line treatment followed by a combination of Cabozantinib with Nivolumab or Pembrolizumab based on expression profile. We found that ARAF (Sorafenib target) showed a significantly higher FC in Asian patients than Caucasian patients, which is consistent with other studies (Table 5). The more significant potential to downregulate ARAF in Asian patients may be associated with improved outcomes. A small retrospective study on Sorafenib has reported that it appears to be better suited for delaying time to progression (TTP) in HCC patients of non-Caucasian and non-African American backgrounds, showing significantly longer TTP in patients of Asian, Hispanic, and other ethnicities.^49^

FGFRL1 was significantly upregulated in Caucasian patients compared with other ethnicity groups; thereby, given its potential to downregulate FGFRL1, Lenvatinib could be a better candidate with superior efficacy as a first-line treatment than Sorafenib for this group. There were no statistically significant results for systemic treatments for Africa American patients. Additionally, gene ontology terms showed that leukocyte mediated immunity and positive regulation of MAPK cascade/processes were the most significant pathways dysregulated in all ethnicity groups during HCC (Figure 4). Thus, additional therapies that disrupt these pathways should be considered.

### The Influence of Etiology in Ethnic Disparities of HCC

The transcriptional landscape of HCC stratified by ethnicity may be a consequence of prevalence of etiologies within each demographic. However, the genetic data available in the TCGA database lacks the necessary sample size to control for both ethnicity and etiology. An example in one US study found that the prevalence of NAFLD has been reported highest in Hispanic patients, followed by Caucasians, and lowest in African Americans.^37^ There was little data in this study to ascertain NAFLD prevalence in Asian patients. In addition, a study conducted by the National Health and Nutrition Survey estimated that the prevalence of chronic HCV is lowest in Caucasians at 1.5%, intermediate in Hispanics at 2.1%, and highest in African Americans at 3.2% (for the population assessed), while prevalence varied between 3%-6% for Asian American patients depending on location in the US.^50^

In addition to evaluating the prevalence of specific etiologies that drives hepatocarcinogenesis, the influence of specific viral genotypes should also be considered. For example, HBV is most common among Asian Americans and non-Hispanic black Americans. While an estimated 0.3% of the country is reported to have chronic Hepatitis B, 70% of cases are reported to be imported from individuals immigrating into the US.^48^ This suggests that the genotypes of HBV vary widely within the US and may reflect distinctive genetic landscape associated with HCC, even within the same etiology. If heterogeneity is prevalent between etiologies, this must be considered when developing biomarkers and treatment for HCC. Neglecting these factors could contribute to cancer health disparities.

### Sample Size - A Limiting Factor Beyond Liver Cancer

The low sample size of African Americans in this dataset was a limiting factor in thoroughly assessing the transcriptional landscape of HCC concerning the geographic-origin of patients. This challenge is not unique to liver cancer, but rather an ethical quagmire within the medical community in which ethnic groups of African, Asian, Hispanic, Alaskan, American Indian, Hawaiian, and Pacific Island descent are routinely underrepresented in genomic sequencing studies and biorepositories.^47^ In the NIH TCGA database, 77% of patients (n = 5729) are of Caucasian ethnicity, which means that large scale genomic assessments diminish deviations in disease pathogenesis between patients of different ethnicities. The small sample size of patients of non-Caucasian ethnicities may correlate to public distrust in the medical community. To fully understand HCC, patient engagement with individuals of varying backgrounds must be sought to appropriately assess, diagnose and treat HCC.

## Conclusion

The development of biomarkers and therapeutics for HCC must consider the reasons for genomic distinctions between patients of different ethnicities. The implications of this preliminary research must be evaluated further to support these claims, especially given that the sample size of African American early stage HCC samples on the TCGA contains only 8 samples. An expansion should assess a specific etiology of HCC with regard to patient ethnicity to determine if etiology contributes to distinct transcriptional patterns during HCC. Patient engagement is crucial to initiate these goals. While conducting appropriate research studies will not solve cancer health disparities, these efforts are a sufficient place to start.

## Acknowledgment

We thank Dr. Bram Lutton, Dr. Timothy McCaffrey and Kate Lacey for critical review of this manuscript.

## Abbreviations

HBV: hepatitis B virus
HCC: hepatocellular carcinoma
HCV: hepatitis C virus
NAFLD: Nonalcoholic fatty liver disease
NASH: Nonalcoholic steatohepatitis
LIHC: Liver Hepatocellular Carcinoma
PNPLA3: polymorphisms in adiponutrin 3
GCKR: Glucokinase Regulator
TCGA: The Cancer Genome Atlas
RNA-Seq: RNA-sequencing
CGC: Cancer Genomics Cloud
AJCC: American Joint Committee on Cancer
SPP1: osteopontin
GOLM1: Golgi Membrane Protein
MDK: midkine
DKK1: Dickkopf-1
GPC3: glycoprotein-3
F2: coagulation factor II
DCPS: Decapping Enzyme Scavenger
FUCA1: alpha-1-fucosidase
AFP: alpha fetoprotein

## Notes

**Potential conflict of interest:** Nothing to report.

**Financial Support:** This work was supported by AGED Diagnostics.

The Genotype-Tissue Expression (GTEx) Project was supported by the <u>Common Fund</u> of the Office of the Director of the National Institutes of Health, and by NCI, NHGRI, NHLBI, NIDA, NIMH, and NINDS. The data used for the analyses described in this manuscript were obtained from: the GTEx Portal on 09/04/2020 and/or dbGaP accession number <u>phs000424.vN.pN</u> on 09/04/2020.

### Competing Interest Statement

AGED Diagnostics.

