## Supplemental Figures and Tables for "Transcriptional Landscape of Hepatocellular Carcinoma Reveals that Patient Ethnic-Origin Influences Patterns of Expression"

**SUPPLEMENTARY FIGURE 1.** RNA-Seq bioinformatics pipeline

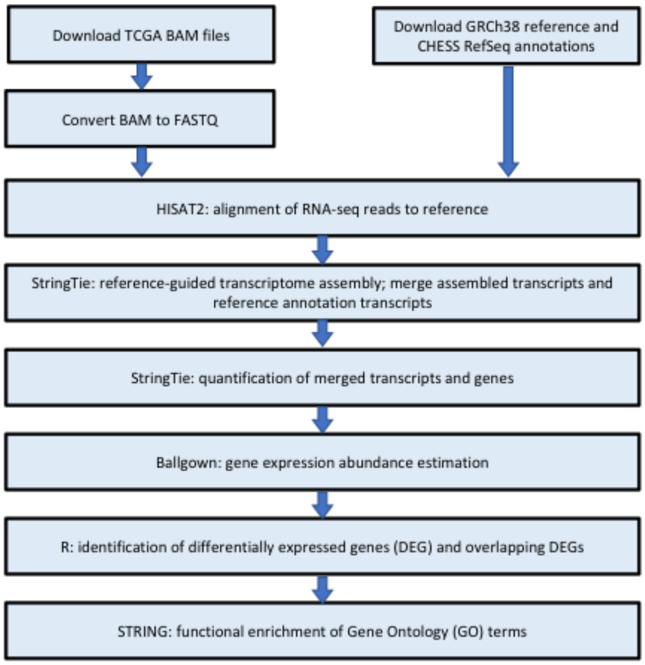


**SUPPLEMENTARY FIGURE 2**. Stage I HCC transcripts among and between each group

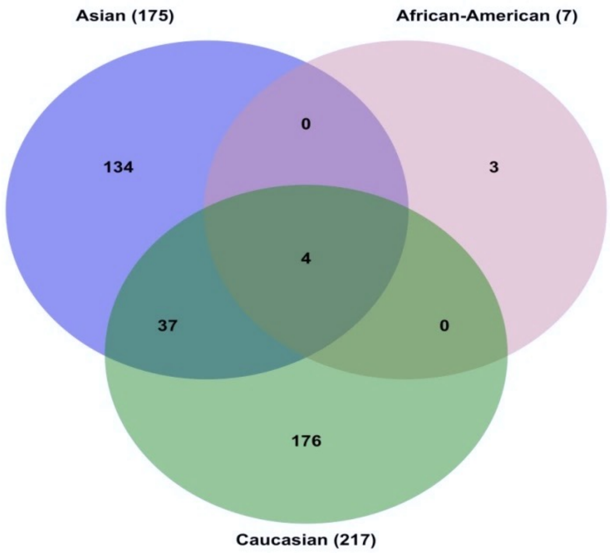

**SUPPLEMENTARY FIGURE 3**. Top differentially expressed gene within each ethnicity group

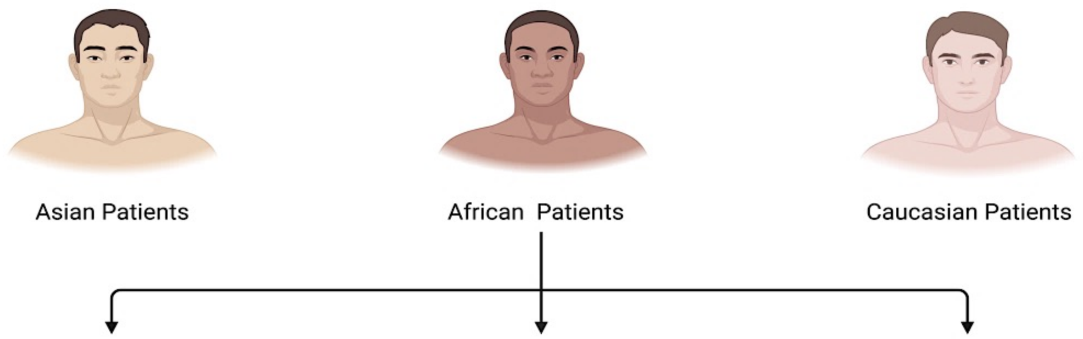


| Genes | Q-value | logFC |
|---|---|---|
| *HAMP* | 0 | 6.17 |
| *FCN2* | 0 | 4.11 |
| *CD5L* | 0 | 2.67 |
| *GDF2* | 0 | 3.00 |
| *GNE** | 6.90E-13 | 2.55 |
| *CD4* | 8.35E-11 | 3.67 |
| *IGFBP3* | 3.37E-10 | 3.12 |
| *HBA2* | 7.51E-10 | 3.20 |
| *SLC38A2** | 1.38E-09 | 4.27 |
| *MARCO* | 1.68E-09 | 4.48 |

| Genes | Q-Value | logFC |
|---|---|---|
| WRNIP1* | 8.17E-06 | 3.55 |
| GDF2 | 1.38E-03 | 2.52 |
| COLEC10 | 8.62E-03 | 2.83 |
| FCN3 | 3.92E-03 | 3.92 |
| FCN2 | 3.97E-03 | 1.75 |
| MYD88* | 3.97E-03 | 1.68 |
| RDH16* | 4.18E-03 | 2.28 |

| Genes | Q-Value | logFC |
|---|---|---|
| FCN2 | 0 | 3.12 |
| MARCO | 3.40E-12 | 3.64 |
| CLEC4G | 8.93E-12 | 3.63 |
| CXCL14 | 2.87E-09 | 2.21 |
| GDF2 | 4.18E-09 | 2.38 |
| CFP | 2.50E-06 | 2.25 |
| CD5L | 5.14E-06 | 3.17 |
| FCN3 | 5.23E-06 | 2.08 |
| HAMP | 4.75E-05 | 2.33 |
| MT1F* | 4.08E-05 | 4.04 |

**SUPPLEMENTAL FIGURE 4**. Liver-specific transcript expression patterns in Stage I HCC in each geographic-origin group

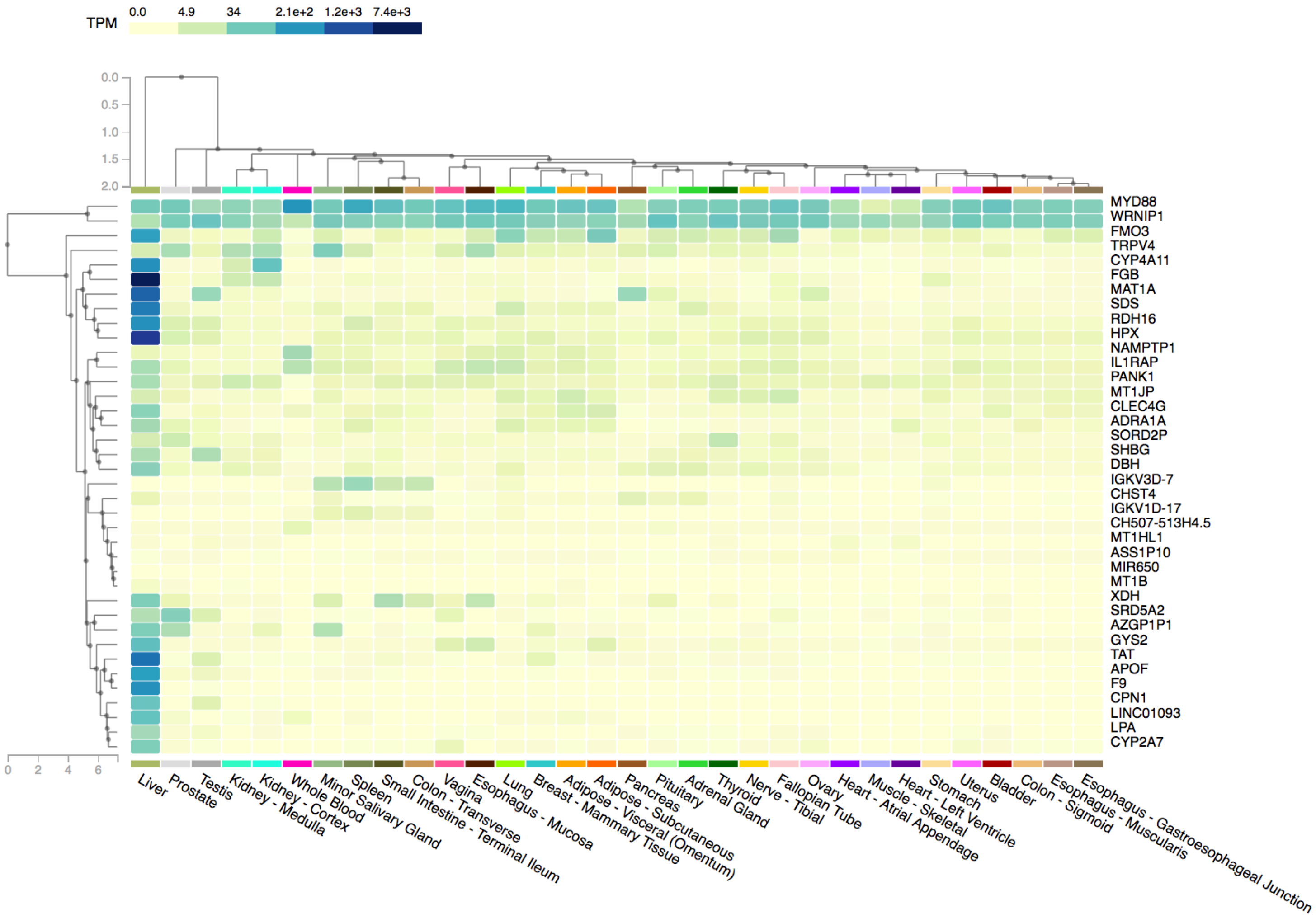

**SUPPLEMENTAL TABLE 1**. Uniquely expressed transcripts in each ethnicity group.

---

**Asian Downregulated Genes**: *UQCRB, VPS28, LBHD1, ITIH3, ZYX, MRPL24, GAS5, CREBL2, TXNDC12, ZNHIT1, CALR, POLRMT, PSMB4.*

**African American Downregulated**: *None.*

**Caucasian Downregulated Genes**: *RBP7, PLVAP, NF2, SLC25A1, NDUFA2, PPIB, BCAT2, TIMM44, AKT1S1, PSMB5, NKAP, ATP6V1F, DRAP1, HULC, PLOD3, TM7SF2, SELENBP1, EBNA1BP2, ATP5J2, MEA1, APOLD1, ACOX2, FKBP11, DYNLRB1, CRELD2, RPA3, ESM1.*

**Asian Upregulated**: *GNE, SLC38A2, SDS, LSR, AZGP1P1, ADAR, CPN1, APH1A, SORD2P, RPL39P3, CERS2, RNF11, MBD3, SLC25A28, ETS2, TDP2, DBH, AKR1A1, ADH1C, STK40, CES2, GADD45G, CAP1, LRPPRC, DNTTIP2, DNAJ, XDH, IDI1, EHHADH, KLKB1, FTCD, EIF5, CFI, INMT, MST1L, C1R, CSRP1, CDHR5, MFAP4, SLC25A20, EPB41L5, LDHD, PUF60, DCTD, MS4A6A, HAO2, PLIN2, BAX, ASS1, DMGDH, GHR, MS4A7, RAD23B, SOCS3, AKT1, SRD5A2, LPA, TXNIP, CEBPZ, ARSD, KIF1C, BTBD1, NAMPTP1, SEC31A, TAT, APOF, APOA1, TRAPPC3, CXCL2, NAPSB, DUSP5, CEACAM1, RPRD1B, SLC37A4, F9, RND3, TMEM123, HPX, MRPL9, GLOD4, AMOTL2, PRKAR1A, RND1, NR4A1, ADAMTSL2, LDLR, FMO3, ITGA9, TLN1, ZSCAN25, PGM1, GANAB, ANPEP, TNFRSF1A, RAMP3, PLXNB2, OXT, SQSTM1, PRSS8, PROS1, CAT, SPIDR, HDLBP, UBTF, ATF5, LAPTM5, ASS1P10, DOK4, YTHDF2, IFI6, ZNF410, STAP2, UBA1, SRRT, PANK1, CNDP1, NRBP1, BCAS2, RHBDD2, FGB, HLA-DRB1.*

***African American Upregulated***: *WRNIP1, MYD88 and RDH16.*

**Caucasian Upregulated**: *CLEC4G, CXCL14, GSTT1, MT1H, MT1G, MT1F, MT1HL1, LINC01093, LOC107984936, MT1M, DUSP2, PTH1R, FOSB, ZFP36L1, CCL3, PLSCR4, LOC105378356, EGR1, MT1B, IGHV5-51, IGHA1, CCL4, ADRA1A, JUN, TACSTD2, IGF1, EPHA2, AK3, GNMT, MASP1, MT1L, MT1JP, CHST4, IL1RAP, TRIB1, LOC107985462, SFRP5, GYS2, CYR61, JUNB, SERPINE1, C5AR1, CYP4A11, LOC107987279, IGKV3D-7, TRPV4, PTGIS, LOC105376156, ALB, ATP6V1C1, IGKV3D-15, FAM180A, EMILIN1, RNA18S5, TTC36, IGLJ3, IGKV3-20, IGHA2, CH507-513H4.5, KCNN2, RASGEF1B, IGLC2, SLC39A5, BCHE, IGKV3D-20, IL1B, IGLV3-25, IGHG1, POLR1E, TMEM45A, ST6GAL1, IGKV3D-11, SHBG, CYP1A1, MT2P1, CCL2, SERPINA1, MT1X, CCL14, IGKV3-15, VNN1, HSD11B1, FGL2, FNDC4, GPR183, RGS1, CD69, CYP2A7, CFTR, LOC107985919, IGKV1D-17, MIR650, CD163, C8orf4, C1QC, IGKV4-1, HCLS1, IGLL5, AGTR1, MT2A, PRODH2, IGLV6-57, NUDC, NDRG2, DPT, KRT19, THBS1, C8A, SRGN, IGLC6, IGKV1-5, DHRS4-AS1, IGLV2-11, IDH1, IGLV2-14, IGLV1-44, IGLV2-23, RNA5-8S5, JCHAIN, SLC2A3, IGKV3-11, SLC6A1, NAT2, SLC25A47, VKORC1, IGKV1D-12, IGKV1D-39, RGN, IGLV2-33, IGHV3-30, IGKV2-24, SSB, IGKV1D-8, IGLV2-8, IGHV3-33, GPD1, KLF10, FAH, IGHG3, PGLYRP2, DUSP6, IGKV2-30, IGLV3-19, CXCR4, RPL18, IGKV1-39, IGLV2-18, IGLV3-21, MAT1A.*

---

**SUPPLEMENTAL TABLE 1**.

| **Ethnicity** | **Asian** | **African American** | **Caucasian** |
|---|---|---|---|
| Total Patients | n = 158 | n = 17 | n=153 |
| Male | 124 | 14 | 86 |
| Female | 34 | 3 | 67 |
| Vital Status | | | |
| Alive | 115 | 11 | 88 |
| Deceased | 43 | 6 | 65 |
| AFP at Procurement | | | |
| <400 ng/mL | 90 | 9 | 87 |
| >400 ng/mL | 33 | 2 | 23 |
| NA | 35 | 6 | 43 |
| Vascular Invasion | | | |
| Macro | 4 | 3 | 10 |
| Micro | 43 | | 36 |
| None | 83 | 11 | 85 |
| NA | 28 | 3 | 22 |
| Stage | | | |
| I | 80 | 8 | 68 |
| II | 35 | 4 | 37 |
| III | 40 | 2 | 32 |
| IV | 1 | - | 2 |
| NA | 2 | 3 | 14 |
| Liver Fibrosis | | | |
| No Fibrosis | 9 | 5 | 48 |
| Portal Fibrosis | 14 | 1 | 14 |
| Fibrous Speta | 18 | 1 | 8 |
| Nodular Formation | 3 | 2 | 3 |
| Established Cirrhosis | 39 | 1 | 25 |
| NA | 75 | 7 | 55 |
| Ablation Therapy | | | |
| Yes | 4 | - | 6 |
| No | 108 | 8 | 87 |

| | | | |
|---|---|---|---|
| NA | 46 | 9 | 60 |
| Etiology | | | |
| HBV | 69 | 2 | 2 |
| Alcohol Consumption | 24 | 1 | 35 |
| No Known Risk Factor | 25 | 1 | 48 |
| NAFLD | 1 | 1 | 9 |
| HCV | 5 | 8 | 15 |
| N/A | 13 | 1 | 15 |
| Two or more factors | 21 | 3 | 24 |
| Hemochromatosis | 0 | - | 5 |

**Figure and Table Legends**

**SUPPLEMENTAL FIGURE 1**. RNA-seq bioinformatics pipeline. BAM files were converted to FASTQ using the SAMtools FASTQ app provided by the CGC and submitted to HISAT2 for alignment to GRCh38 reference and CHESS RefSeq annotations. The StringTie application performed transcriptome assembly and quantified merged transcripts and genes. Gene expression abundance estimation was determined by the Ballgown application, and output was downloaded locally for additional processing using R (Version 4.0.1). Post-hoc analysis of Ballgown output included the overlapping of differentially expressed genes and transcripts, and coercion into formatting appropriate for STRING and GO-term analysis on Cytoscape (Version 3.8.0).

**SUPPLEMENTAL FIGURE 2.** Stage I HCC transcripts between and among each ethnicity. Venn Diagram depicting the overlap in shared transcripts and differences in transcripts between each ethnicity group during Stage I HCC.

**SUPPLEMENTAL FIGURE 3**. Top differentially expressed genes within each ethnicity group, sorted by lowest P-value. Cutoff criteria of absolute value FC > 2.0, p-value < 0.001 and q-value <0.05.

**SUPPLEMENTAL FIGURE 4.** Healthy liver-specific transcript expression patterns in Stage I HCC in each ethnicity group. Using the GTEx Database Multiple Gene Query Assessment and the transcripts dysregulated exclusively in each ethnic-origin group, heatmap was constructed to assess liver specificity. Map is clustered by tissue. Legend (top left) depicts extent of transcript expression in transcripts per million (TPM). Transcripts that were found greater than 67 TMP in multiple other healthy tissues were filtered out of dataset. Transcripts assessed for Caucasian Stage I HCC included IL1RAP, TRPV4, CLEC4G, CYP4A11, IGKV3D-7, MT1JP, ADRA1A, LINC01093, GYS2, CHST4, MT1HL1, MT1B, SHBG, CYP2A7, CH507-513H4.5, MIR650, IGKV1D-17 and MAT1A. Transcripts assessed for Asian Stage I HCC included FGB, ASS1P10, PANK1, HPX, FMO3, NAMPTP1, SRD5A2, LPA, TAT, APOF, F9, SORD2P, DBH, XDH, AZGP1P1, CPN1 and SDS. Transcripts assessed for African American included WRNIP1, MYD88 and RDH16.

**SUPPLEMENTAL TABLE 1**. Uniquely expressed transcripts in each ethnicity group. Uniquely expressed genes are defined as a group of genes that were up- or down-regulated in one specific ethnicity group with the absence of significant changes in the other two groups assessed. African American patients had 3 upregulated transcripts and no downregulated transcripts. Asian patients had 121 upregulated transcripts and 13 downregsulated transcripts. Caucasian patients had 149 upregulated transcripts and 27 downregulated transcripts.

**SUPPLEMENTAL TABLE 2.** Expanded Patient Sample Demographics. HCC Demographics including gender, vital status, AFP at procurement, vascular invasion, AJCC Neoplasm Disease Stage Classification, fibrosis score, ablation therapy, and etiology. Abbreviation: alpha fetoprotein, AFP. nonalcoholic fatty liver disease, NAFLD, Hepatitis C Virus, HCV, Hepatitis B Virus, HBV, American Joint Committee on Cancer (AJCC).
